# Analysis of cholesterol export from endo-lysosomes by Niemann Pick C2 protein using combined fluorescence and X-ray microscopy

**DOI:** 10.1101/462481

**Authors:** Alice Dupont, Frederik W. Lund, Maria Louise V. Jensen, Maria Szomek, Gitte K. Nielsen, Christian W. Heegaard, Peter Guttmann, Stephan Werner, James McNally, Gerd Schneider, Sergey Kapishnikov, Daniel Wüstner

**Author notes:** These authors contributed equally (Alice Dupont, Frederik W. Lund, Maria Louise V. Jensen). Address correspondence to: Daniel Wüstner, Department of Biochemistry and Molecular Biology, University of Southern Denmark, Campusvej 55, DK-5230 Odense M, Denmark.

## Abstract

The Niemann-Pick C2 protein (NPC2) is a sterol transfer protein in late endosomes and lysosomes (LE/LYSs). How its capacity to transport cholesterol between membranes is linked to endo-lysosomal membrane trafficking is not known. Using quantitative fluorescence imaging combined with soft X-ray tomography (SXT); we show that NPC2 mediated sterol efflux is accompanied by large changes in distribution, size and ultrastructure of endocytic organelles. We observed clearance of intra-luminal lipid deposits, a decrease in number of autophagosomes, formation of membrane contact sites (MCSs) to the endoplasmic reticulum and extensive tubulation of LE/LYSs in three-dimensional SXT reconstructions of NPC2 treated human fibroblasts. The cells could recycle the cholesterol analog dehydroergosterol (DHE) from LE/LYSs slowly also in the absence of NPC2 protein but internalized NPC2 synchronized and accelerated this process significantly. Most fluorescent NPC2 was retained in LE/LYSs while DHE was selectively removed from these organelles, at least partially by non-vesicular exchange with other membranes. During sterol efflux LE/LYSs were reallocated to the cell periphery, where they could fuse with newly formed endosomes. Surface shedding of micro-vesicles was found, suggesting a pathway for cellular sterol release. We conclude that NPC2 mediated sterol efflux from LE/LYSs controls membrane traffic through the endo-lysosomal pathway.

## Introduction

Late endosomes and lysosomes (LE/LYSs) play a central role in uptake and processing of nutrients, but these organelles are also increasingly recognized as sensors of the cellular nutrient status ^1^. Cholesterol, an essential cellular lipid ^2^ is delivered to cells mostly via uptake of low density lipoproteins which become processed in LE/LYSs to provide cholesterol for other cellular destinations. Two proteins, NPC1 and NPC2 are critically involved in export of cholesterol from LE/LYSs. According to a well-established model, the small soluble sterol-binding protein NPC2 picks up cholesterol inside LE/LYSs and delivers it to NPC1 for export from these compartments.^3, 4^ Importantly, lack of both, NPC1 and NPC2 leads to Niemann Pick type C (NPC) disease, in which cholesterol and sphingolipids accumulate in LE/LYSs of liver spleen and neuronal cells to a varying degree^5-7^. The tissue-specific accumulation of cholesterol versus various sphingolipid species caused some disagreement of the initially accumulating metabolite in NPC disease ^8^. Both, NPC1 and NPC2 can bind cholesterol and other sterols and NPC2 accelerates sterol exchange between liposomes in a concentration-dependent manner under conditions found in LE/LYSs ^4, 9-14^. Despite this wealth of biochemical in-vitro data, the molecular mechanisms underlying the function of both proteins in cholesterol transport in living cells are not clear.

There is increasing appreciation of the role played by the intracellular positioning of LE/LYSs in the function of these organelles. For example, there is evidence for a direct involvement of lysosome positioning in the nutrient status of the cell^15^. Cholesterol enrichment in LE/LYSs due to defective rab7 GTPase or NPC1 can cause dynein-dependent perinuclear accumulation and reduced mobility of LE/LYSs ^16-19^. In fact, the ion composition of LE/LYSs is tightly regulated and directly affects ligand receptor sorting and cargo degradation ^20^. We found recently, that fluorescent NPC2 mobilizes cholesterol visualized using filipin from NPC2 deficient fibroblasts (NPC2 -/-cells), and that this process is accompanied by increasing mobility and dispersion of LE/LYSs ^21^. Filipin staining, however, does not allow one to distinguish different pools of sterols, as all sterols bearing a free 3’hydroxy group bind filipin ^22^. One can overcome this limitation by using fluorescent analogs of cholesterol, which can be specifically inserted into the plasma membrane (PM) to follow their intracellular trafficking over time using quantitative fluorescence microscopy ^22^. Here, intrinsically fluorescent cholesterol analogs like dehydroergosterol (DHE) should be preferentially used instead of dye-tagged cholesterol analogs, as the latter are often poor mimics of cholesterol in membranes, while the dye moiety can interfere with sterol binding to enzymes and transfer proteins ^23^. DHE and other intrinsically fluorescent sterols do not bear attached fluorescent moieties and can be recognized by various sterol transporters including Oxysterol-binding proteins, StARD3, StARD4, NPC1 and NPC2 ^9, 24-29^. In a recent study, we found that PM-derived DHE and isotope-labeled cholesterol becomes trapped in LE/LYSs in NPC2 -/-cells ^30^. Internalized NPC2 could rescue transport of this sterol between PM and LE/LYSs, partly by increasing non-vesicular DHE export from the storage organelles. The molecular mechanisms underlying re-mobilization of PM-derived sterol from LE/LYSs by NPC2 are poorly understood. Also, it is not clear how this process is orchestrated with the spatial distribution of these organelles and whether NPC2 mediated sterol efflux affects membrane trafficking through the lysosomal pathway in general.

In order to address these questions, we have performed a detailed quantitative imaging study to monitor NPC2 mediated efflux of PM-derived sterol from LE/LYSs of NPC2-/-fibroblasts. We show that DHE trapped in LE/LYSs can slowly efflux from patient fibroblasts also in the absence of functional NPC2. Internalized NPC2 accelerates and synchronizes this efflux process and co-localizes extensively with DHE during efflux. Sterol efflux was accompanied by reallocation of LE/LYSs from the perinuclear region to the cell periphery and extensive tubulation of these organelles, as shown by both, live-cell fluorescence microscopy and soft X-ray tomography (SXT). The latter is an emerging technique, which provides 3D images of fully hydrated cryo-frozen cells with a 3D resolution down to 36 nm (Rayleigh) ^31, 32^. SXT is therefore extremely useful to study the ultrastructure and spatial arrangement of LE/LYSs. We use SXT to provide ultrastructural information on endosomes, their size and spatial relationship to other organelles during cholesterol efflux. We found that NPC2 has a profound impact on the ultrastructure, location and dynamics of LE/LYSs. Due to its excellent preservation and detection of membranes, SXT allowed us also to detect delicate cellular structures, like shedding micro-vesicles at the PM, which might be implicated in sterol release. Our study combines different optical imaging modalities to provide a mechanistic link between the observed in vitro function of NPC2 and the cellular phenotype of NPC2 disease.

## Results

### Kinetics of sterol efflux and ultrastructure of endo-lysosomes are controlled by NPC2

We studied first the dynamics of cellular egress of accumulated sterol in disease cells after removal of sterol sources from the culture medium. When NPC2-/-fibroblasts were incubated in the continuous presence of DHE/BSA for 48 h, the fraction of fluorescent sterol in the LE/LYSs rises to about 50% of total cell-associated DHE ^30^. NPC2 removed this excess sterol to levels found in fibroblasts from healthy subjects (Fig. 1A and B). When these cells were subsequently incubated in medium having LPDS but no sterol or NPC2, some DHE was released from the cell after a prolonged delay phase (Fig. 1B, black symbols). When Alexa488-NPC2 was present during the incubation in LPDS (Protocol A, Fig. S1), the delay was much shorter (approx. 8 h) followed by a pronounced drop of DHE in the lysosomal storage compartments (Fig. 1B, dark red symbols). At the end of the 96 h incubation period, the fraction of intracellular DHE was ~0.38 in disease cells not treated with Alexa488-NPC2, while it was ~0.28 in disease cells treated with NPC2. Under exactly the same conditions, the intracellular DHE fraction was ~0.18 in control fibroblasts incubated in LPDS without Alexa488-NPC2 (Fig. 1B, blue symbol). Remobilization of trapped sterol after removal of the sterol source was slow in cells incubated just in medium with LPDS compared to cells incubated in the presence of Alexa488-NPC2. Sterol efflux could not be followed for longer than four days without compromising cell function.

**Figure 1.**
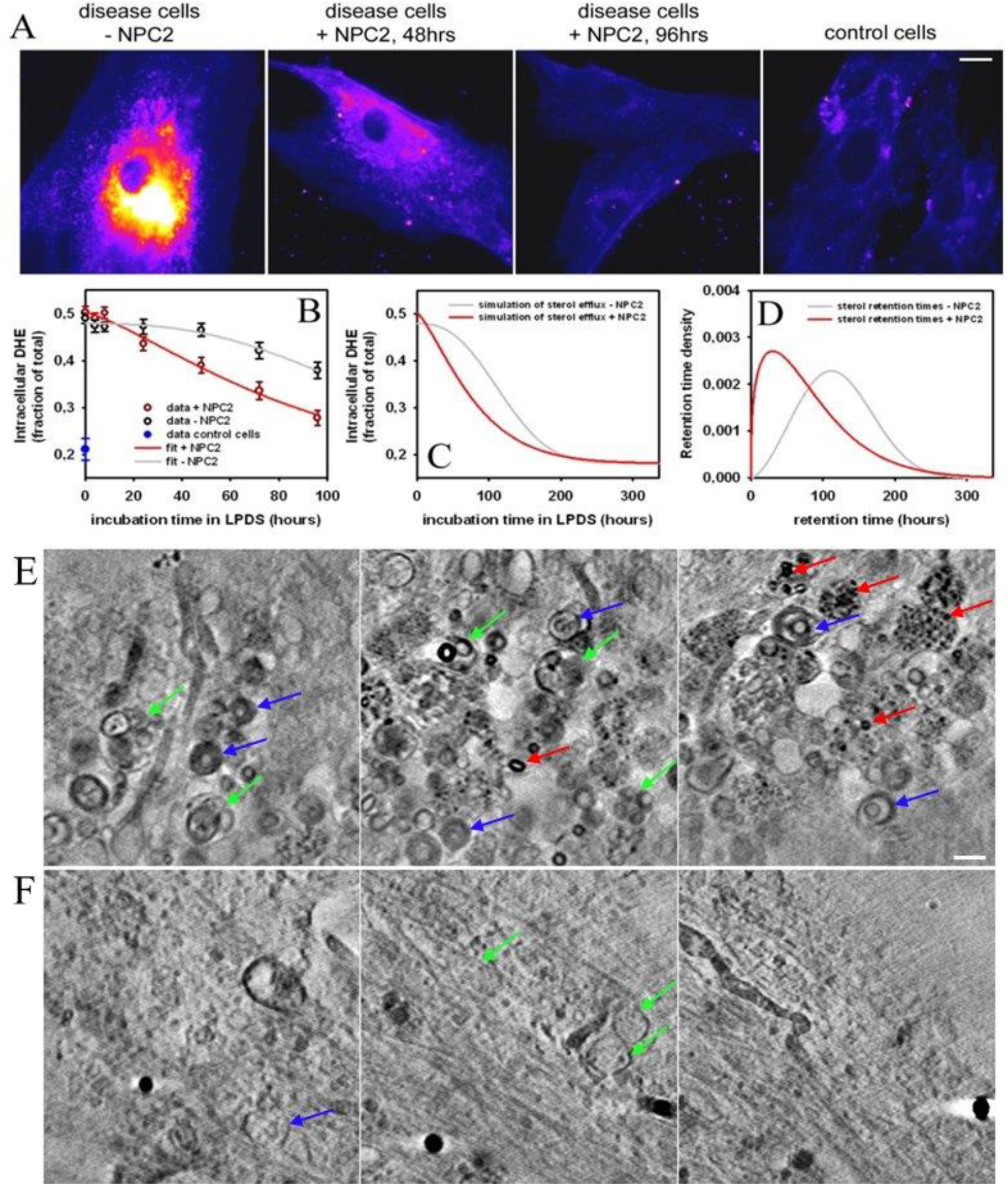
Efflux of DHE from disease fibroblasts in the presence and absence of NPC2. NPC2-/-cells were loaded with DHE for 48 h, washed and incubated in medium containing LPDS, either in the absence (-NPC2) or presence of 100 nM Alexa488-NPC2 (+ NPC2) for the indicated times (Protocol A, Fig. S1). A, intensity images of DHE pseudo-colored with high intensities in yellow/white and low intensities in blue. Most right panel shows control fibroblasts loaded with DHE/BSA for 48 h for comparison. Bar, 10 μm. B-D, quantification of DHE in LE/LYSs of cells incubated without (black symbols) or with NPC2 protein (red symbols) is shown in (B). The data was fitted to a Weibull function (grey and red lines in B). Simulation of this function with the fit parameters shows that DHE efflux takes place without NPC2 protein (C) but NPC2 strongly reduces the sterol retention time (D). E, F selected sum projections of SXT reconstructions along the optical axis covering a thickness of 0.24 μm each for NPC2-/-cells incubated in the absence (E) or presence of 100 nM NPC2 (F). Multi-vesicular bodies (green arrows), autophagosomes (blue arrows) and LE/LYSs with dark lipid deposits (red arrows) are indicated. Bar, 0.5 μm.

The observed delay in sterol efflux in disease cells in the absence of NPC2 indicates initial hindrance and temporal heterogeneity of sterol release. Such temporal delays are often observed in drug dissolution studies from heterogeneous, i.e., not well-mixed compartments ^33^. To test this and to quantify sterol efflux rates, we made use of a stochastic compartment model in which one calculates the probability that a particle ‘survives’ in a compartment until a given time. We used a compartment model based on the Weibull distribution and fitted this model to the time-dependent fractional fluorescence of DHE in LE/LYSs of NPC2-/-cells incubated in the presence or absence of NPC2 (see Appendix). The data was gathered from three biological replicates with six to eight technical replicates (i.e. imaged fields with 2-4 cells each) for each time point. We found that the efflux kinetics of DHE in NPC2-/- fibroblasts in the presence and in the absence of Alexa488-NPC2 can be exactly fitted by the Weibull model function (red and grey line in Fig. 1B). For this analysis, the DHE fraction in the LE/LYSs of control cells (blue symbol in Fig. 1B) was used as background term to equation (2), see Appendix. Accordingly, the amplitude of the decay, *S*_0_, describes the sterol efflux back to normal levels found in control cells under the same loading conditions. The estimated time constants from fitting the model to the data were τ = 86.96 h for cells incubated in the presence of NPC2 and τ = 135.14 h in the absence of NPC2. The values for the stretching parameter, which describes the width of the decay function, were μ = 1.324 and μ = 2.577 for cells incubated in LPDS in the presence and absence of Alexa488-NPC2, respectively. Using these estimated parameters, we can predict that sterol efflux proceeds beyond four days and will be complete after about ten days irrespective of the presence of Alexa488-NPC2 (Fig. 1C). This information could not be inferred from the experiment alone, since the disease fibroblasts showed signs of stress when cultured in LPDS without NPC2 for longer than five to six days (not shown). The stochastic efflux of sterol is based on an intracellular retention time distribution (or probability density function; *pdf*) of sterol molecules ^33^. In contrast to a simple exponential-type efflux, where efflux would be equally likely throughout the experiment, the likelihood for a sterol molecule to leave the cell according to the Weibull model is different at the start and at the end of the efflux experiment (see Appendix), which is often defined as a ‘memory’ of the *pdf* ^33^. Using the estimated parameters from the fit of equation (2) to the sterol efflux data, we can calculate the *pdf* for the sterol efflux in the presence and absence of Alexa488-NPC2 according to equation (3) and find that the *pdf* for sterol efflux is narrowed and skewed to shorter times in the presence of NPC2 compared to that for sterol efflux from cell just incubated in LPDS (Fig. 1D). In addition, the rate coefficient is growing steadily over time for the efflux in NPC2-treated cells, while it levels of in cells incubated in LPDS without NPC2 (Fig. S2). From that, we can conclude that internalized fluorescent NPC2 significantly reduces the retention time and in parallel synchronizes the sterol efflux process within and between the cells.

To gain a better understanding of how NPC2 may function at the sub-cellular level, we used soft X-ray tomography (SXT) of NPC2-/-cells prior to and after incubation with NPC2. This optical microscopy technique attains increased resolution compared to classical light microscopy by using X-rays with λ=2.3-4.4 nm (equivalent to an energy of 280-540 eV) ^34^. In that range, water does not absorb, while inner shell electrons of carbon, being very abundant in cells and tissues, get excited leading to absorption contrast. In SXT, the specimen is cryo-frozen and tilted while projection images are taken over a large range of tilt angles. Contrast is generated according to Lambert-Beer’s law, meaning that the extent of ‘darkness’ in an image is linearly related to the amount of absorbed light. Especially, the carbon-rich lipids appear dark and give high contrast ^31, 35^. In addition, the cryo-preservation process used in SXT maintains the cell in a near-native state with full hydration, preventing artefacts of altered endosome morphology, often observed in EM after chemical fixation ^36^. Thus, SXT is a quantitative imaging technique, and after reconstruction of the 3D volume covering an entire cell thickness of about 10 μm from the projection series, ultrastructural details not resolvable by classical light microscopy can be determined. We have imaged NPC2-deficient fibroblasts by SXT, prior to and after ‘treatment’ with 100 or 200 nM reconstituted NPC2. In the absence of NPC2, many multi-lamellar or –vesicular endosomes were found by SXT (Fig. 1E, green arrows). Many of the large vesicles harbored dark internal structures with a diameter of ~50 nm, suggesting that they contain remaining lipid-rich intraluminal structures (Fig. 1E; red arrows and Fig. S3). In addition, vesicles with the characteristic double-membrane shape of autophagosomes were observed (Fig. 1E: blue arrows). These results suggest that NPC2-/- fibroblasts are in a state of starvation despite the lipid storage phenotype. Efflux of DHE in cells incubated with NPC2 was accompanied by a dramatic change in the ultrastructure of LE/LYSs with fewer autophagosomes, disappearance of the dark intra-luminal lipid deposits and generally more transparent membrane structures (Fig. 1F). These results demonstrate that internalized NPC2 has a profound effect on the ultrastructure of endo-lysosomes, suggesting that this protein controls not only sterol export from LE/LYSs but also the participation of these organelles in endocytic trafficking processes in general.

### Sterol export by internalized NPC2 requires membrane traffic between endo-lysosomes

To further investigate the sub-cellular function of NPC2, we used fluorescence microscopy to and determined colocalization of NPC2 (Alexa488-NPC2) with cholesterol (DHE). We found that these two molecules were extensively colocalized in small peripheral vesicles (Fig. 2A) that became increasingly scattered in the course of sterol chase-out. In addition, DHE was sometimes found in elongated vesicles and tubules with Alexa488-NPC2 confined to the drop-shaped end of a tube (Fig. 2A, especially lowest panels named ‘96h’ boxes (1) and (2), respectively). Importantly, the peripheral vesicles and to a smaller extent the perinuclear LE/LYSs containing DHE and Alexa488-NPC2 could be labeled with Rh-dextran delivered to the cells at the end of the incubation with Alexa488-NPC2 (compare first three columns in Fig. 2A; ‘Rh-Dex’), suggesting that remobilized LE/LYSs are fusion-competent for incoming cargo. Thus, these organelles are not a ‘dead end’ of the degradative pathway but highly involved in membrane trafficking processes. The peripheral vesicles containing DHE and Alexa488-NPC2 seem to intermingle also with the early endocytic recycling pathway, since they could be labeled with Alexa647-Tf given to the cells after the long incubation with Alexa488-NPC2 (see fourth column in Fig. 2A; ‘Alexa647-Tf’). In cells, which had been pre-incubated with NPC2 small elongated endosomes were confirmed by SXT in the cell periphery at a distance less than one micron to the PM (Fig. 2B). Elongated endosomes and tubules with a diameter less than 100 nm have been previously described in mouse embryonic fibroblasts during co-uptake of cholera toxin and dextran, suggesting that they are actively involved in membrane traffic with the PM ^37^. SXT also revealed that peripheral LE/LYSs were often found to be enwrapped by transparent membrane sheets, likely resembling the ER. In fact, tubule-forming endosomes with internal lipid deposits surrounded by lamellar membranes of the ER were repeatedly observed (Fig. 3 and Supplementary video 1). Such membrane contact sites (MCSs) will likely assist the tubule formation and thereby endocytic sorting process ^38^, while they also could be places of sterol exchange between LE/LYSs and the ER ^39^. In addition, also autophagosomes with the characteristic double membrane shape were found to be enwrapped by ER-like membranes (Supplementary video 1; green arrows). The number of autophagosomes per 100 μm^2^ was determined from SXT reconstructions of two representative cells for each condition and found to be 6.52 in NPC2+/+ cells, 26.26 in NPC2-/-cells incubated without and 12.58 for NPC2-/- cells incubated with NPC2. Thus, clearance of autophagosomes seems to accompany the sterol efflux mediated by NPC2. We conclude that mobilization of excess sterol from LE/LYSs requires NPC2 and makes LE/LYSs fusion competent, able to relocate to the cell periphery and to form membrane tubules and MCSs with the ER.

**Figure 2.**
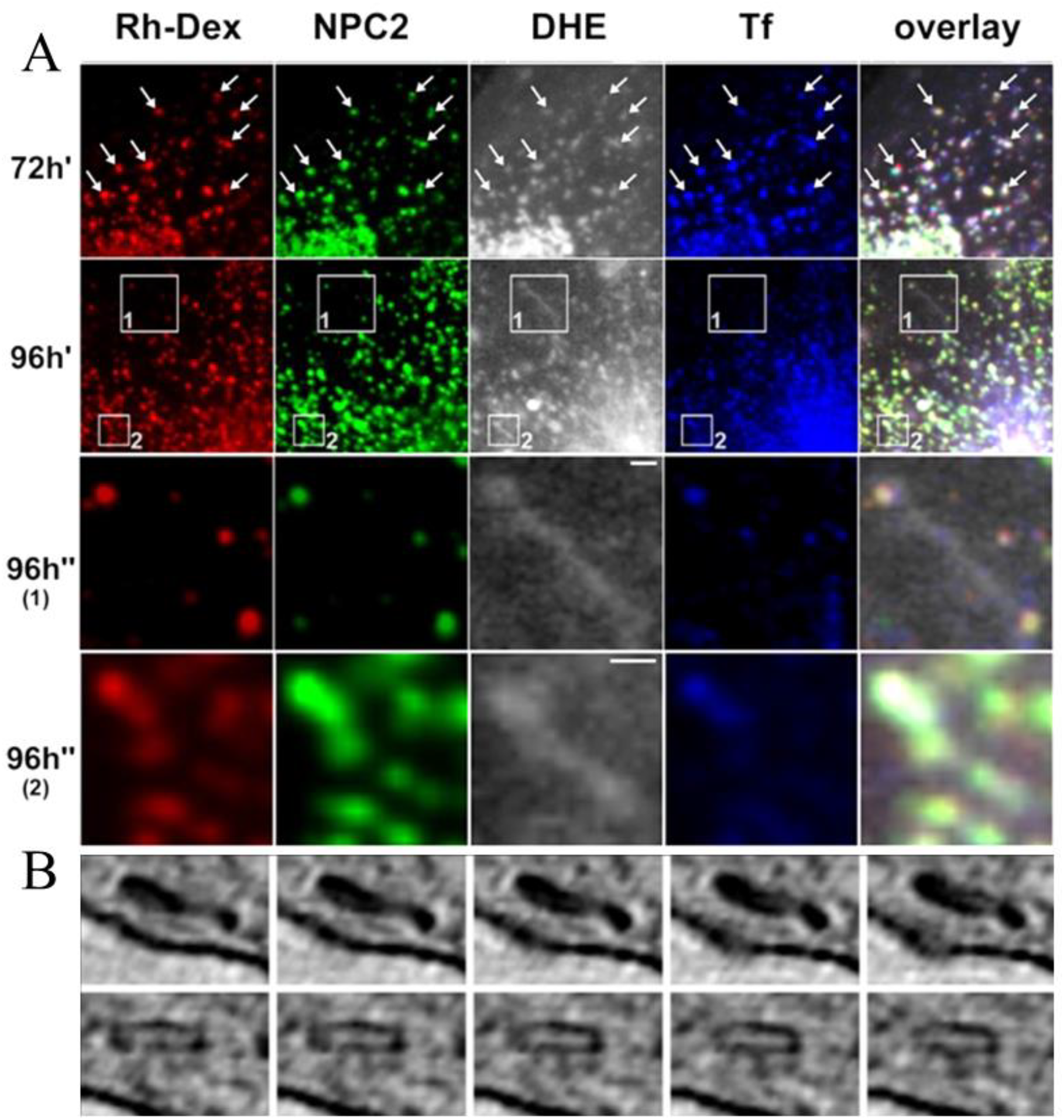
LE/LYSs containing Alexa488-NPC2 co-localize with DHE and endocytic markers. A, NPC2 deficient fibroblasts were incubated with DHE/BSA for 48 h, washed and incubated in culture medium containing LPDS, either in the absence (-NPC2) or presence of 100 nM Alexa488-NPC2 (+ NPC2) for the indicated times (Protocol A, Fig. S1). NPC2-/-cells were co-labeled with Rh-dextran for the last 24 h in the presence of Alexa488-NPC2, and after washing with Alexa647-Tf for 30 min. Arrows indicate co-localization. Bar, 1 μm. B, NPC2-/-fibroblasts were incubated in the presence of LPDS with 100 nM Alexa546-NPC2 for 48 h under sterile conditions, washed, fixed with 4% PFA and kept in buffer before cryo-plunge freezing. Cells were imaged at the TXM at BESSY II, as described in Materials and Methods. A montage of selected frames of a reconstruction of a tomogram series highlighting two examples of tubular vesicles close to the PM (dark curved line) is shown. The width of the panels in B is 1 μm.

**Figure 3.**
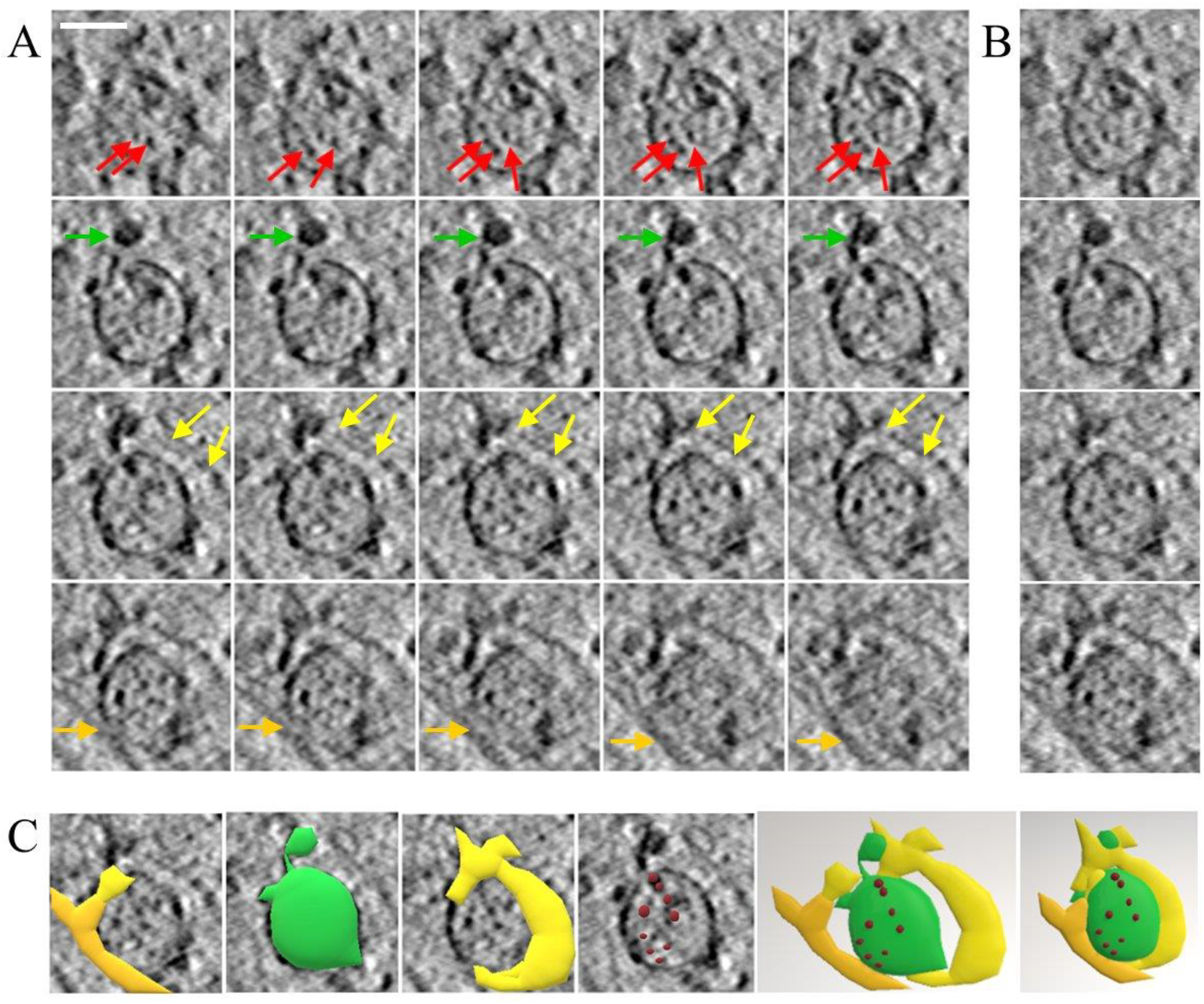
SXT of tubule-forming endo-lysosomes in contact to the ER. NPC2-/-fibroblasts were incubated in the presence of LPDS with 100 nM Alexa546-NPC2 for 48 h under sterile conditions, washed, fixed with 4% PFA and kept in buffer before cryo-plungefreezing. Cells were imaged at the TXM at BESSY II, as described in Materials and Methods. A, montage of selected frames of a reconstruction of a tomogram series showing a multivesicular endosome forming a tubule (green arrows) on the upper left part and a membrane contact site to another organelle, likely the ER at the upper central, right (yellow arrows) and lower central part (orange arrows). This endosome also harbors internal lipid deposits appearing as dark spots (red arrows). B, sum projection of the corresponding 5 frames of each row in panel A along the optical axis. C, shape outline of the observed features showing the endosome in green, the intra-endosomal lipid deposits in red and the enwrapping membranes as they emerge in the 3D-stack in yellow. Bar, 0.5μm.

Since the function and intracellular dynamics of endocytic organelles in general, and of LE/LYSs in particular, are intimately linked to the size of these organelles ^40^, we sought to find how this is affected in NPC2-/-cells. Apart from cargo content, populations of endosomes can be distinguished based on their size, and enlarged late endocytic compartments have been described in various lysosomal storage disorders or upon disruption of the endocytic trafficking machinery ^41, 42^. LEs and LYSs continuously fuse forming larger hybrid organelles, from which smaller new LYSs are re-formed ^43^. Since SXT provides the fully hydrated 3D volume at much higher spatial resolution than obtainable by fluorescence microscopy, we used tomograms reconstructed from SXT data to measure the size of endo-lysosomes in NPC2+/+ and in NPC2-/- cells, incubated in the presence and absence of NPC2. We measured the diameter of all vesicles with approximately round shape at their equatorial plane from tomographic reconstructions in the cell between nucleus and periphery of several cells (see Materials and Methods). The advantage of using SXT compared to electron microscopy (EM) for this is, that one can find the equatorial plane for each endosome separately in SXT, avoiding tedious and error-prone corrections for off-set positions, as needed for analysis of EM sections ^42^. Endo-lysosomes were significantly larger in NPC2-/- cells than in NPC2+/+ cells, and this increased vesicle size could be reverted by treating NPC2-/- cells with the NPC2 protein (Fig. 4). We found a left-skewed size distribution under all conditions, which could be well-described by a log-normal distribution (not shown). The mean vesicle diameter was ≈ 0.5 μm in NPC2+/+ cells and in NPC2-/- cells treated with NPC2, but was close to 0. 7 μm in NPC2-/- cells incubated without NPC2 (Fig. 4A and B). Similar values and size distributions have been determined previously for LE/LYSs by EM ^42^. By SXT, we observed a variety of vesicle forms, as depicted in Fig. 4C. Multi-vesicular endosomes had the largest size of up to ~ 1.5 μm in diameter containing transparent and spherical intraluminal vesicles (ILVs) with a diameter of about 70-100 nm (Fig. 4B and C). By SXT we were able to detect also small vesicles with a diameter centered around 180 nm, which match the size previously described for LEs and LYSs, while larger vesicles might resemble hybrid organelles generated by fusion of both organelle types ^44, 45^. The increased size of endo-lysosomes in NPC2-/- cells incubated in the absence of NPC2 suggests a defect in re-formation of LYSs from hybrid organelles generated by fusion of LEs and LYSs as suggested previously using sucrose loaded LE/LYSs ^46^.

**Figure 4.**
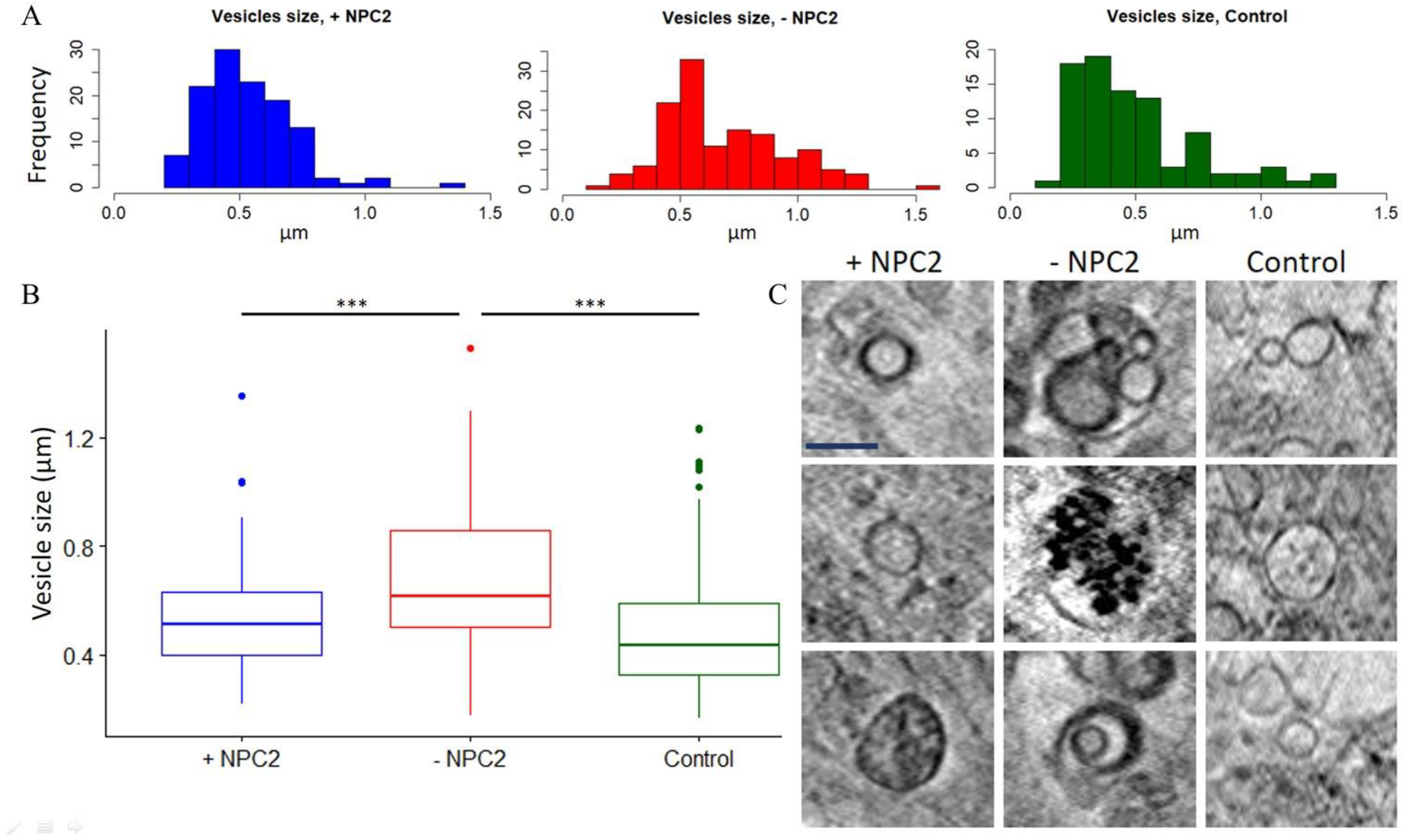
Sterol efflux by internalized NPC2 is accompanied by a reduction of vesicle size. NPC2-deficient fibroblasts were incubated with 100 nM Alexa546-NPC2 in medium containing LPDS for 48 h under sterile conditions, washed, fixed with 4% PFA, kept in buffer before cryo-plunge freezing. Cells were imaged at the TXM at BESSY II, as described in Materials and Methods. Figure 4. A, Histograms of endosome sizes (um) in NPC2 lacking (n=134, N=3), 72 h NPC2 loaded (n=120, N=3) and control fibroblasts (n=86, N=1). B, Boxplot showing the quartiles, minimum, maximum, outliers and median for NPC2 lacking (mean= 0.6923164, median=0.6181722), 72 h NPC2 loaded (mean = 0.533629, median = 0.512) and control (mean=0.4979454, median=0.43375) fibroblasts. The datasets of the endosome sizes were transformed to log normal distributions for which ANOVA and Tukey HSD tests were performed. These tests showed significant differences between NPC2 lacking and NPC2 rescued cells (p=3.6∙10^−6^), and between NPC2 lacking and control fibroblasts (p=1.0∙10^−8^), while no significant difference were found between the endosome sizes of control and NPC2 rescued cells (p=5.2636∙10^−2^). C, Representative vesicles types in NPC2 lacking, NPC2 rescued and control fibroblasts. Size estimation of endo-lysosomes vere derived as mean of horizontal (yellow line) and vertical (white line) measurements. Bar, 0.5 μm.

### Intraluminal NPC2 decreases the DHE intensity in LE/LYSs and allocates them to the PM

To determine, whether fluorescent NPC2 acts from inside LE/LYSs when stimulating sterol efflux, we loaded cells first with DHE for 24 h and subsequently allowed for uptake of 200 nM Alexa546-NPC2 for another 24 h. Such cells were next washed and incubated further in medium with LPDS during which sterol efflux was monitored and quantified (Protocol B, Fig. S1). We developed an image analysis protocol to detect and segment not only LE/LYSs vesicles in the red channel (i.e. Alexa546-NPC2) but also in the UV channel (i.e. DHE) (Fig. S4). This was necessary, as DHE vesicles have to be identified in the presence of DHE fluorescence in the PM which gives a bright background distribution (see Supplemental information, Fig. S4 to S7). When quantifying fluorescence on a vesicle basis using this new image analysis protocol, we observed a continuous drop in DHE intensity per endo-lysosomal vesicle, while the intensity of fluorescent NPC2 decreased only a little (Fig. 5). These results demonstrate that NPC2 indeed acts from inside LE/LYSs where it mediates the selective removal of sterol from the organelles. This same approach has been used previously to show that cyclodextrin removes sterols from inside LE/LYSs in NPC1 and NPC2 defective fibroblasts including those used here ^47^.

**Figure 5.**
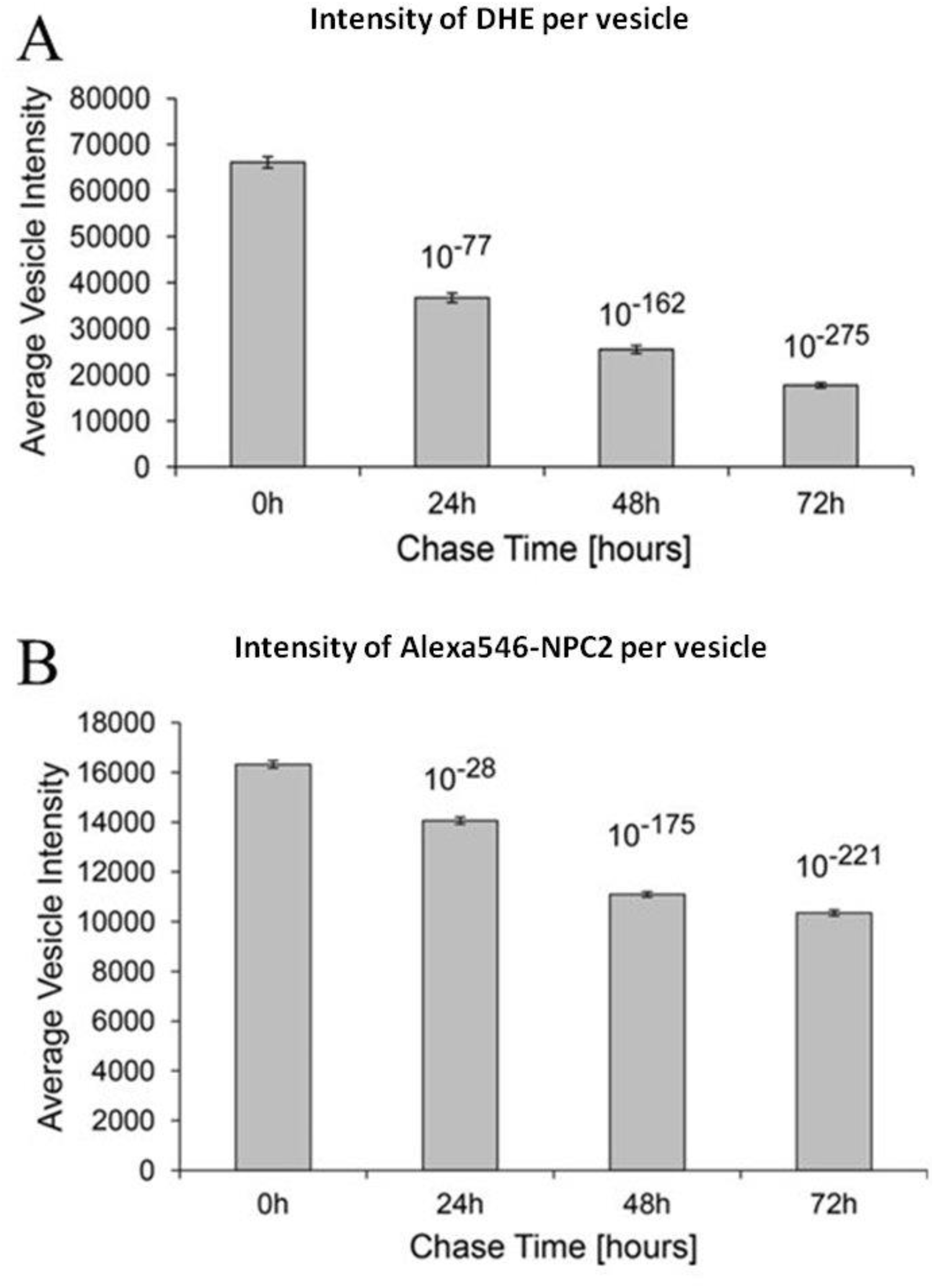
DHE is preferentially released from individual LE/LYSs during sterol efflux. NPC2 deficient fibroblasts were incubated with DHE/BSA for 24 h, washed, incubated with 100 nM Alexa546-NPC2 in culture medium containing LPDS for 24 h, followed by a chase in the absence of NPC2 for up to 72 h (Protocol B, Fig. S1). The average intensities of the DHE vesicles (A) was determined from ~7000 vesicles per time point while the average vesicle intensity for Alexa546-NPC2 (B) was determined from ~15000 vesicles. Data is shown as mean ± standard error. Note, that the intensity values are in arbitrary units, such that the numerical values cannot be directly compared between both panels.

Using spatial statistical analysis of the determined endosome positions^21^, we observed that LE/LYSs enriched in DHE were initially clustered but reallocated to the cell periphery during NPC2 mediated sterol efflux (Fig.6A-D). This result is in accordance with earlier studies reporting a direct effect of the endo-lysosomal cholesterol content on the spatial location of these organelles 16,17. We determined co-localization of DHE and Alexa546-NPC2 in an object-based manner, i.e., separately for each detected LE/LYS in both fluorescence channels throughout the whole cell; we found that ~50-55% of the DHE containing vesicles were co-localized with NPC2 containing vesicles at all time points of efflux (Fig. 6E and F). On the other hand, the percentage of NPC2 containing vesicles which also have significant amounts of DHE increased from ~22% at 0 hours to ~30% at 72 hours. Thus, Alexa546-NPC2 is also found in vesicles with low sterol content, while more than half of the DHE-rich vesicles found throughout all cells contain fluorescent NPC2. DHE and Alexa546-NPC2 co-localized increasingly in vesicles confined to a narrow region of about 3 μm from the PM in the course of sterol efflux (insets in Fig. 7A and quantifications in Fig. 7B-C). Thus, sterol-rich, NPC2 containing vesicles continue to reallocate to the PM also after removal of NPC2 from the culture medium. At the same time, the number of vesicles containing DHE remained constant (Fig. S8A, B). To explain that observation, one could assume that sterol release takes place from individual vesicles by lysosomal exocytosis accompanied by compensating sterol endocytosis generating new vesicles containing some DHE derived from the PM by endocytosis. In this model, the overall number of sterol-containing vesicles could remain constant, such as we observed. However, such a scenario would also predict that we find an increasing number of vesicles close to the PM containing DHE but not Alexa546-NPC2, since NPC2 cannot be reinternalized once in the medium. This, however, was not observed, since the number of vesicles containing Alexa546-NPC2 remained also constant (Fig. S8C, D). Thus, complete fusion of sterol-rich LE/LYSs with the PM for sterol release is an unlikely scenario in our experiments.

**Figure 6.**
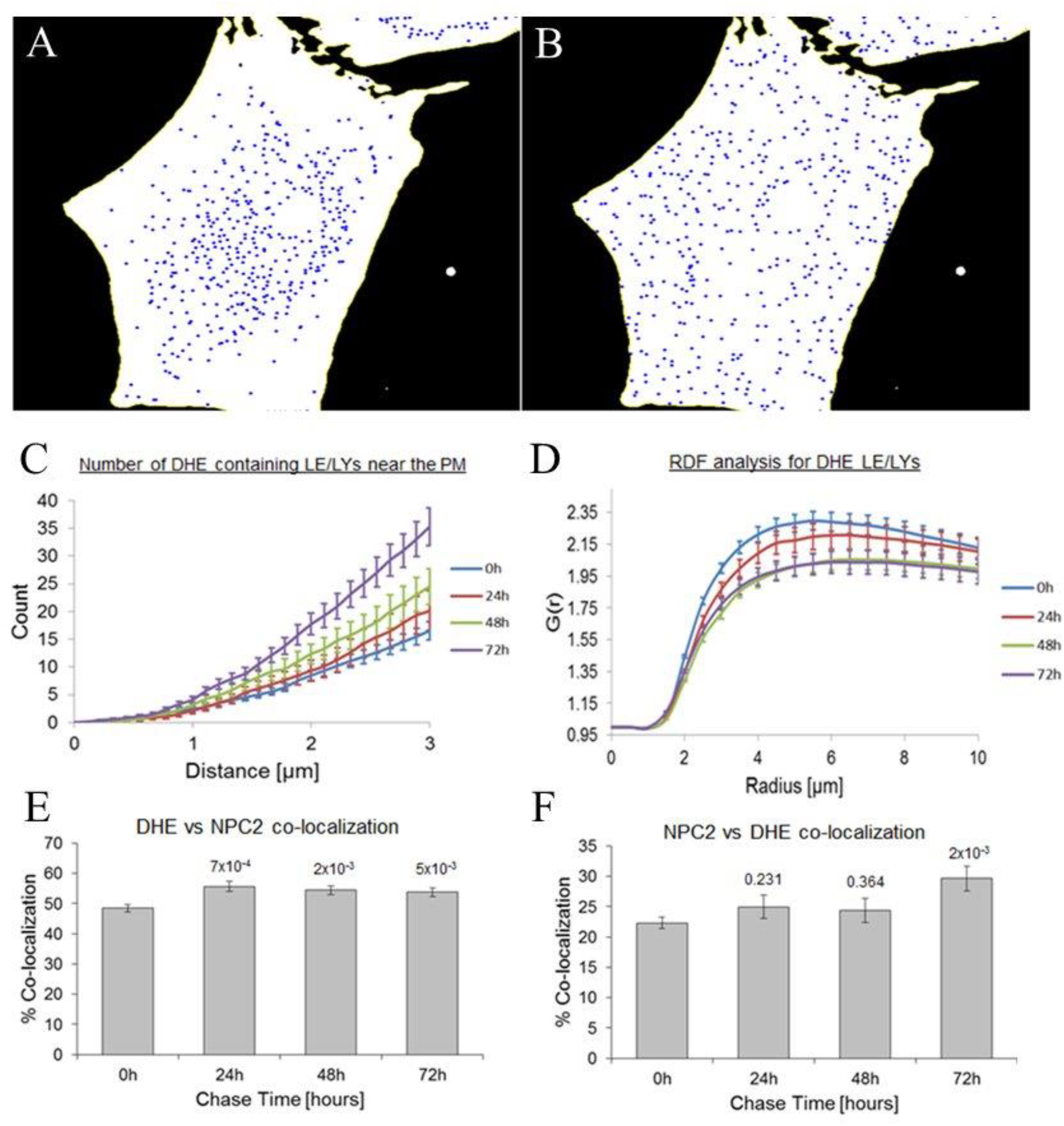
Spatial pattern and co-localization analysis of DHE and NPC2. NPC2 deficient fibroblasts were incubated with DHE/BSA for 24 h, washed, incubated with 100 nM Alexa546-NPC2 in culture medium containing LPDS for 24 h, followed by a chase in the absence of NPC2 for up to 72 h (Protocol B, Fig. S1). DHE and Alexa546-NPC2 were imaged on a wide field microscope, and images were analyzed using SpatTrack (see Materials and Methods). A, B, centroid positions of DHE containing vesicles (A) and the same number of randomly placed particles (B) within the cell geometry, as derived by segmenting DHE stained cell images. C, D, number of DHE containing vesicles as a function of distance to PM (C) or of inter-particle radius (D, RDF) for different chase times after internalization of Alexa546-NPC2. E, F, particle-based co-localization analysis of DHE and Alexa546-NPC2 containing vesicles for the same chase times.

**Figure 7.**
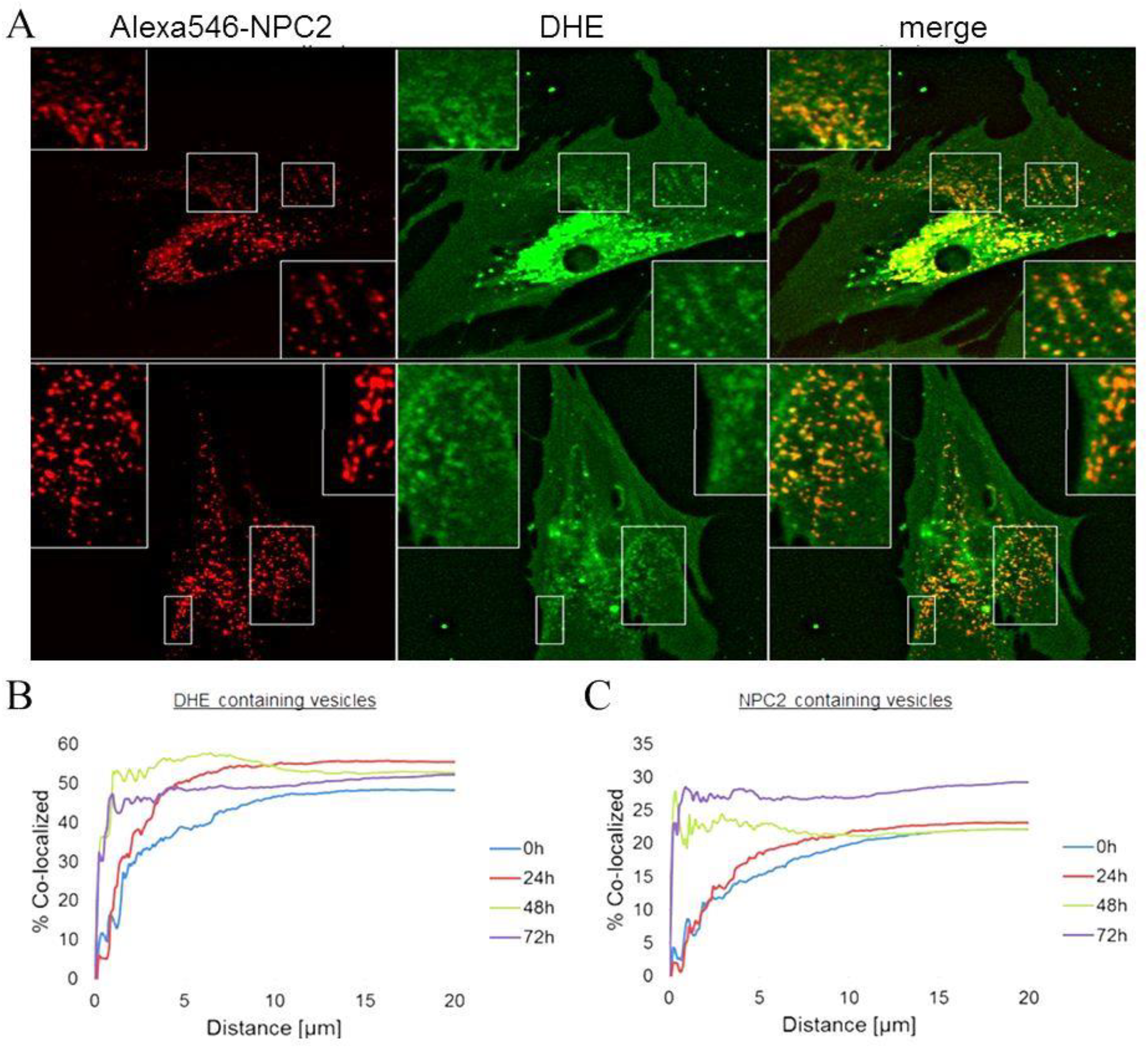
DHE co-localizes with Alexa546-NPC2 in peripheral LE/LYSs during sterol efflux. NPC2 deficient fibroblasts were incubated with DHE/BSA for 24 h, washed, incubated with 100 nM Alexa546-NPC2 in culture medium containing LPDS for 24 h, followed by a chase in the absence of NPC2 for up to 72 h (Protocol B, Fig. S1). A, example images after chase for 24 h (upper row) and 72 h (lower row) in medium containing LPDS but no NPC2 protein. The boxes and insets indicate regions, in which clear co-localization of DHE and Alexa546-NPC2, especially close to the PM is found. B, C, particle-based co-localization analysis for DHE vs. Alexa546-NPC2 (B) and vice versa (C) as function of distance to the PM, found in segmented cell images based on DHE’s fluorescence in the PM. Colors indicate mean co-localization quantified from three experiments with 4-8 image fields with 1-3 cells per field and 250-400 vesicles per cell for 0h (blue), 24 h (red), 48h (green) and 72h (violet) chase in the absence of NPC2 in the medium.

### Direct observation of vesicle shedding at the PM by soft X-ray tomography

NPC2 is required for maintenance of normal sterol transport between LE/LYSs and the PM ^4^, and internalization of NPC2 rescues the abnormal sterol storage phenotype in LE/LYSs of NPC2-/- cells (Fig. 1, 5 and 7). We verified in independent experiments, that the vesicular structures found by SXT overlap with the fluorescence signal of Alexa546-NPC2, though structural details cannot be resolved in our setup (Fig. S9). Thus, most of the vesicles observed by SXT in NPC2-/- cells treated with fluorescent NPC2 should contain this protein. But how is sterol released from cells once it arrives at the PM? One possibility is the direct excretion of intraluminal lipid content from endosomes upon their transient fusion with the PM, and we found indeed ultrastructural evidence for such events by SXT (Fig. S10). Another possibility is the exo-vesiculation of the PM followed by release of PM derived vesicles – also called ectosomes - into the medium. Shedding of micro-vesicles from the PM has been proposed to be a potential mechanism by which cells communicate, and such vesicles were found to be enriched in cholesterol ^48, 49^. However, micro-vesicles are very small and unless enriched by a suitably fluorescent marker, hard to detect by fluorescence microscopy. Furthermore, simply detecting small vesicles in the culture medium would be by itself not sufficient, as exosomes derived from exocytosis of ILVs of multi-vesicular endosomes would have a similar size and morphology as ectosomes.

We therefore searched for capturing vesicle formation events directly at the PM fibroblasts by SXT. We found examples of vesiculation of the PM in control fibroblasts (Fig. 8A) and in NPC2-/- fibroblasts treated with Alexa546-NPC2 (Fig. 8B), but not in disease cells incubated in the absence of NPC2. Forming vesicles were between 100 and 200 nm in diameter, which is a characteristic value for shedding ectosomes ^49^. During vesicle budding the neck of the forming vesicle was either very narrow and of high contrast with a pear-shaped vesicle attached (Fig. 8A) or broad and of rather low contrast compared to the forming vesicle (Fig. 8C). Narrow necks with dilating vesicles on the exoplasmic site could indicate that some energy barrier caused by shear stress of the protrusion must be overcome, for example by relief of PM-cytoskeleton attachment^50^. Exoplasmic shedding of micro-vesicles looked different from classical endo- or exocytosis, which we also repeatedly observed in the SXT reconstructions (Fig. 8B). Next to the side of micro-vesicle shedding we observed cytoplasmic small vesicles (Fig. 8B). Such vesicles have been proposed to deliver material for shedding to the PM ^49^. A reconstruction of the 3D stack of the vesicle formation event of Fig. 8C is shown after segmentation in Fig. 8D. Together, shedding of small vesicles or ectosomes from the PM into the exoplasmic medium likely resembles another route of selective sterol release during NPC2 mediated sterol efflux.

**Figure 8.**
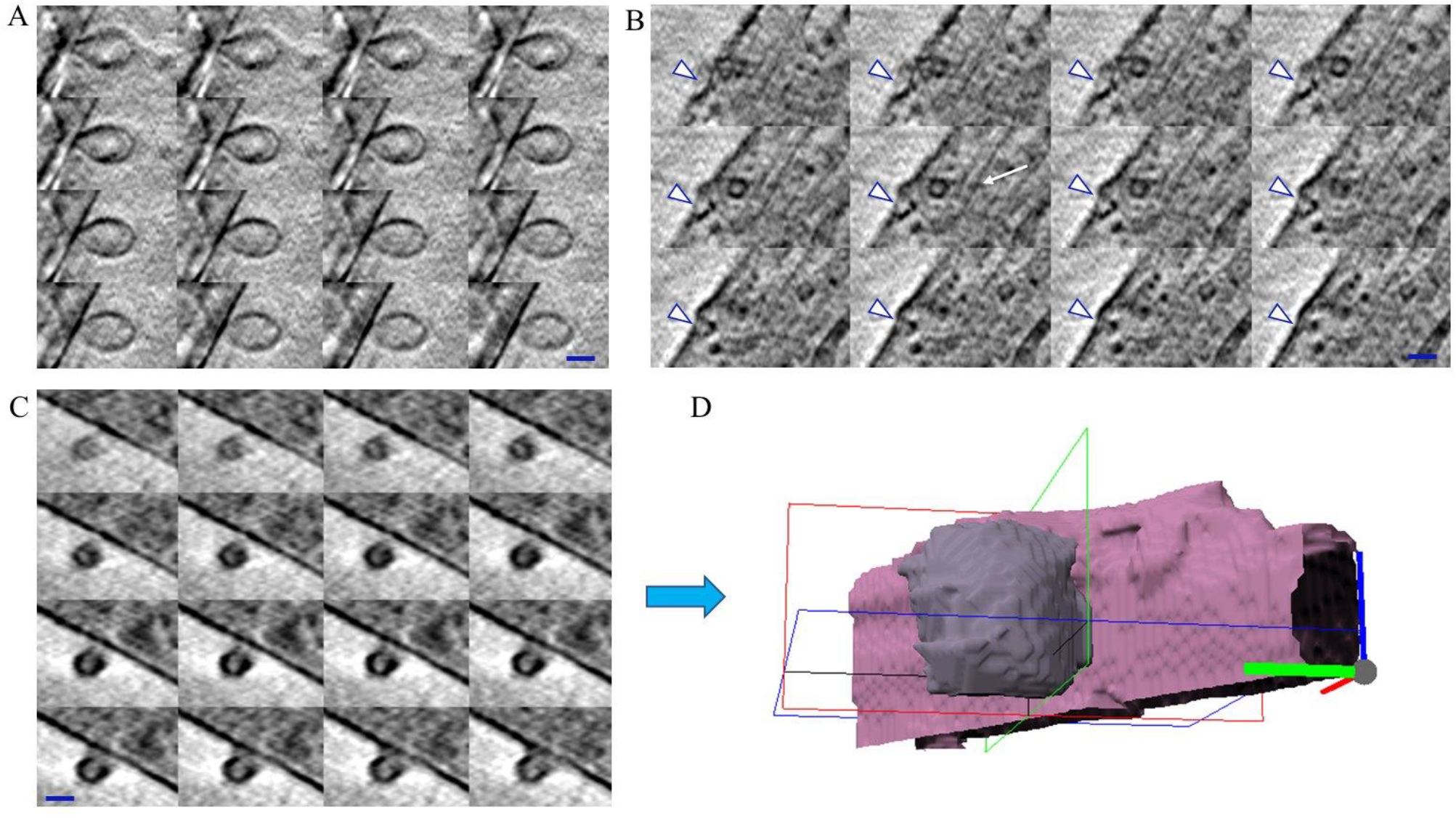
SXT of vesicle formation at the exoplasmic and cytoplasmic sites of the PM. Control (A) or disease fibroblasts (B-D) were washed, fixed with 4% PFA, kept in buffer before cryo-plunge freezing and imaged at the TXM at BESSY II, as described in Materials and Methods. NPC2-/- cells were pre-incubated in the presence of LPDS with 100 nM Alexa546-NPC2 for 48 h under sterile conditions. A-C, montage of selected frames of a reconstruction of a tomogram series showing the ultrastructure of micro-vesicles at the extracellular site of the PM (A, C) as well as an inward (endocytosis) or outward (exocytosis) budding vesicle (B). D, surface rendering of the shedding microvesicle at the exoplasmic site from panel C. Bar, 0.15 μm.

## Discussion

Cellular cholesterol homeostasis depends critically on sterol flux through LE/LYSs, but the pathways and molecular mechanisms of sterol trafficking between these organelles and other membranes are poorly defined ^51, 52^. In Niemann-Pick type C2 disease, cholesterol and other lipids build up due to lack of a small sterol-transfer protein, NPC2, in the lumen of LE/LYSs. Here, we have used NPC2-deficient fibroblasts as a model system to study the spatiotemporal orchestration of export of the close cholesterol analog DHE from LE/LYSs by quantitative fluorescence microscopy and to determine the ultrastructure of lipid carriers during efflux by SXT.

Using this combined microscopy approach, we found that NPC2-/- cells accumulate large amounts of DHE and harbor aberrant multi-vesicular endosomes and an increased number of autophagosomes in the peri-nuclear region (Fig. 1 and 4). Increased autophagy despite excess accumulation of cholesterol has been previously described for NPC2 deficient cells ^53^. Removal of the external sterol source by incubating cells in medium without NPC2 caused release of some of the excess DHE, and fitting of the efflux time course to a stochastic compartment model allowed us to conclude that this process eventually could remove the majority of trapped DHE (Fig. 1A-D). Similar results have been reported in NPC1-deficient fibroblasts suggesting that the capacity of disease cells to get rid of endo-lysosomally trapped sterol is strongly compromised but not totally blocked ^7, 54, 55^. It is likely that albumin in the LPDS containing medium acts as sterol acceptor in the absence of NPC2 in our experiments ^56^. Internalized bovine NPC2 removes the excess sterol from LE/LYSs over several days, and this process takes place with increasing speed and synchronization, as concluded from the stochastic model fitted to the kinetic efflux data (Fig. 1 and S2). The increasing pace of sterol efflux in the presence of NPC2 was characterized by an increasing rate coefficient for the stochastic compartment model (Fig. S2) and was accompanied by an augmenting co-localization of DHE with Alexa488-NPC2 (see Fig. 2). Internalized fluorescent NPC2 co-localizes not only with DHE but also with incoming endocytic cargo, as shown for Rh-dextran and Alexa647-Tf (Fig. 2).

Fluorescent NPC2 and DHE co-localize increasingly close to the PM during efflux, and such vesicles are frequently elongated or tubulated and enwrapped by ER membranes (Fig. 2 and 3). Sterol efflux continues after removing NPC2 from the medium, showing that the protein indeed acts from inside LE/LYSs to remobilize the PM-derived trapped DHE (Fig. 5). Supporting that notion, we measured a selective release of DHE over fluorescent NPC2 on a per-vesicle basis suggesting that NPC2 is largely retained in LE/LYSs during sterol efflux. Previously, we reported that LE/LYSs containing Alexa546-NPC2 became increasingly mobile and moved preferentially by active transport in the course of lysosomal cholesterol mobilization ^21^. Increased motility, repositioning and tubulation of LE/LYSs have been recently reported as being intimately linked to restoration of LE/LYS function in lysosomal storage diseases ^57, 58^. Also, NPC2 has been previously implicated in membrane fission during reformation of LYSs from endo-lysosomal hybrid organelles, and this process is known to be accompanied by extensive tubulation of the involved organelles ^46^. Tubulated endo-lysosomes are characteristic for termination of autophagy in response to starvation, when clearance of autophagosomes has ended and LYSs reform ^59^.

What are the intracellular trafficking events underlying NPC2 mediated sterol efflux from cells? NPC2 must act from inside LE/LYSs as we found selective sterol release from LE/LYSs over days after removing the protein from the medium (Fig. 5 to 7). Based on our data and the given in-vitro function of NPC2 as a sterol transfer protein, we propose a model in which non-vesicular sterol exchange between ILVs and the limiting membrane of LE/LYSs are enhanced by NPC2 which is followed by sterol export to the ER and other organelles via MCSs (Fig. 9; Step 1). NPC2 inside LE/LYSs could supply sterol from ILVs to the limiting membrane of LE/LYSs in such a sterol transfer process, which is supported by a several-fold enhancement of bi-directional exchange of cholesterol and DHE between liposomes by NPC2 ^12, 14, 60^. Supply of sterol from ILVs to the limiting membrane of LE/LYSs is also strongly supported by our recent study, in which we found by fluorescence recovery after photobleaching (FRAP) an approximately 2.7-fold higher amplitude of fluorescence recovery of DHE in NPC2-/- cells incubated in the presence of NPC2 compared to those left without NPC2 ^30^. We observed in additional FRAP experiments that after photobleaching, DHE’s fluorescence can recover in discrete, almost immobile vesicles, independent of the vesicle position in the bleached region (Fig. S11 and Supplementary video 2). Recovery of DHE fluorescence across the whole ROI measured as radial intensity profile was on average constant (not shown) in contrast to diffusion-limited recovery, which would give a Gaussian profile evolving in time ^61, 62^. Thus, FRAP of DHE is in line with fast cytoplasmic sterol diffusion and rate-limiting sterol exchange (binding/unbinding) with LE/LYSs. Thus, while in most perinuclear endo-lysosomes DHE’s fluorescence recovers only a little during the experiment, rapid sterol exchange from some vesicles is possible. This underlines the importance of a sterol export system on the cytosolic site, and the extensive contact between LE/LYSs and ER membranes observed by SXT provides a possible path by which sterol could be transferred to other organelles after being made available for export by intraluminal NPC2 (Fig. 3 and 9). The relief in sterol load reinstalls the fusion competence of LE/LYSs resulting in clearance of autophagosomes and further sterol-rich LE/LYSs followed by reformation of LYSs from hybrid organelles (Fig. 9; Step 2 and 3). Indeed, NPC2 has been implicated in the latter process by controlling membrane fission from enlarged hybrid organelles ^46^. Supporting such a process, we observed tubule formation from LE/LYSs (Fig.3) and a reduction of endo-lysosome size in NPC2-/- cells after internalization of NPC2 (Fig. 4). After some normalization of membrane sterol content, the LE/LYSs should regain their ability to exchange rab proteins, like rab7 and its effector RILP, thereby initiating their centripetal transport and tubule formation, e.g. via kinesin motors ^17^ ^18, 63^. In fact, kinesin motors have been shown to mediate tubule formation mainly from large LE/LYSs as this required less pulling force compared to nanotube formation from smaller vesicles in in-vitro experiments ^64^. Tubule-containing endocytic compartments are also often observed during autophagic reformation of LYSs ^65^, and one can speculate that this process takes place during NPC2 mediated cholesterol efflux as well, as we observed tubulation of LE/LYSs and clearance of autophagic vacuoles in NPC2-/- cells treated with NPC2 (see Fig. 2, 3 and 9). We observed previously an increased contribution of active transport to the mobility of NPC2 containing vesicles over the course of cholesterol efflux ^21^. This active transport will shift the steady state distribution of LE/LYSs towards the PM thereby accelerating further sterol exchange and cellular release (Fig. 9; Step 4) ^21^. The initially hindered but accelerating sterol efflux kinetics (Fig. 1) could well be a consequence of such regained mobility, fusion competence and sterol exchange capability of LE/LYSs. Once allocated to the PM, further non-vesicular sterol exchange might take place between LE/LYSs containing NPC2 and the PM until eventual sterol gradients are balanced ^66^. This could restore the membrane fusion machinery on the endosomes which is known to be inhibited by excessive cholesterol load ^67^. We found that the vesicle associated DHE intensity decreases much more than that of fluorescent NPC2 (see Fig. 5 and 7). In addition, the number of LE/LYSs remains stable in the course of sterol efflux (Fig. S8). Thus, lysosomal exocytosis with complete fusion of LE/LYSs is not a likely mechanism for sterol release. Instead, we suggest that sterol is released from LE/LYSs by their transient fusion with the PM in a kiss-and-run fashion, eventually secreting their ILVs as exosomes (Fig. 9; Step 5). This scenario is supported by our SXT experiments, where we found multi-vesicular endosomes in close contact or even merging with the PM (Fig. S10). ILVs are the precursors of exosomes and are rich in cholesterol, ceramide and LBPA^68, 69^. Secretion of exosomes has been observed to ameliorate the cholesterol storage phenotype in glia cells deficient for functional NPC1 or NPC2 ^70, 71^. Kiss-and-run mediated sterol efflux has also recently been suggested for LDL-derived fluorescent BODIPY-cholesterol in A431 cells ^72^. Close apposition of some endosomes with the PM, as observed by fluorescence (Fig. 7) and SXT (Fig. S10) supports such a scenario. In parallel to that mechanism, DHE could exchange in a non-vesicular way between LE/LYSs and the PM followed by shedding of small vesicles or ectosomes to the extracellular space (Fig. 9; Step 6). Indeed, we observed membrane vesiculation events by SXT in direct support of such a mechanism (Fig. 8) ^48^.

**Figure 9.**
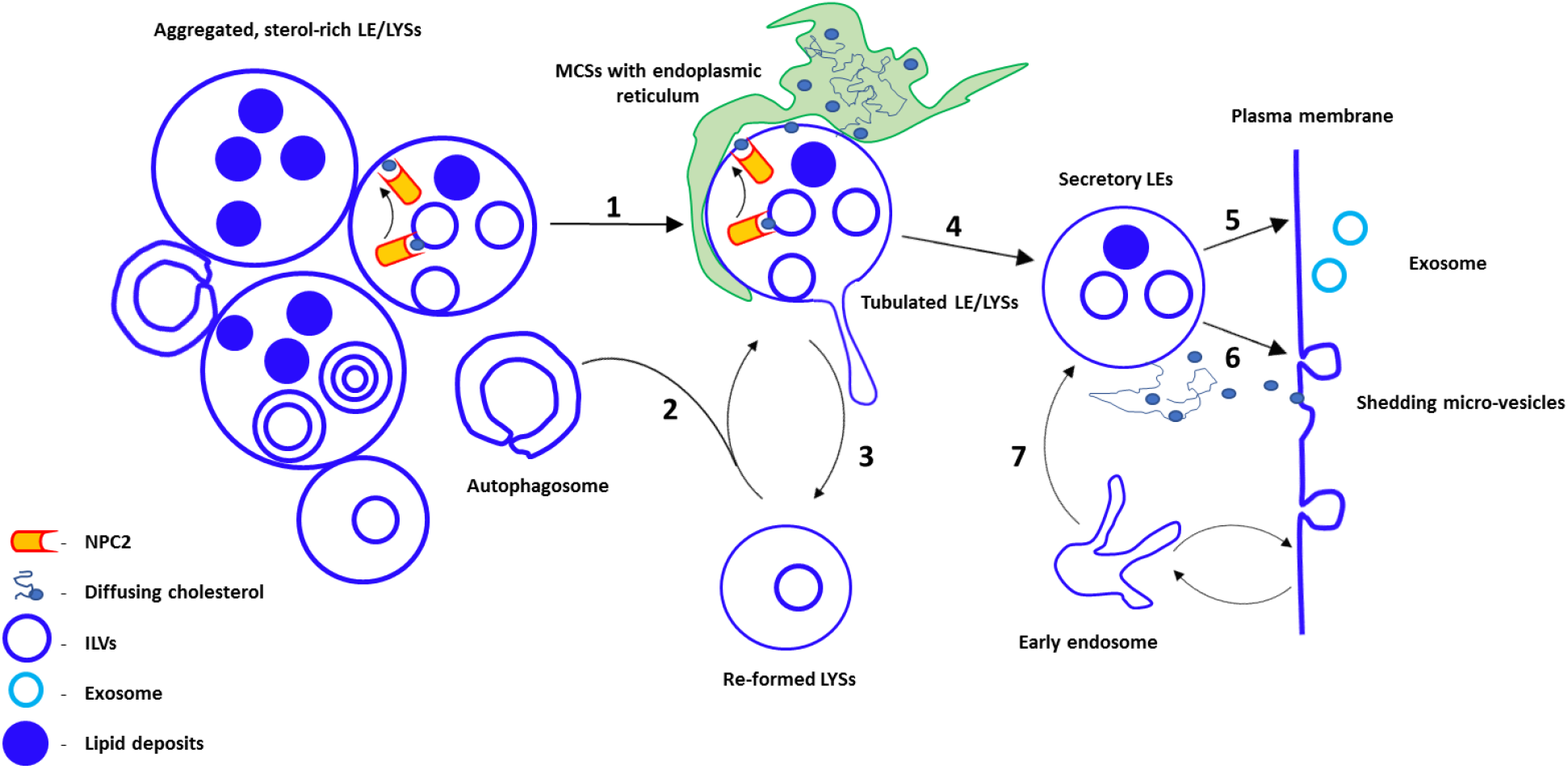
Model describing NPC2’s function in sterol export from LE/LYSs. Prior to addition of NPC2 LE/LYSs with many internal lipid deposits (dark blue spheres) and ILVs as well as autophagosomes accumulate in the perinuclear region. Internalized NPC2 (orange) shuttles sterol monomers (small light blue spheres) from ILVs to the limiting membrane of LEs for export to other organelles. This causes dispersion and tubulation of LEs which also form MCSs to the ER (green; Step 1). Upon relief of excessive cholesterol load, LEs fuse with LYSs and autophagosomes resulting in further clearance of lipid deposits (Step 2). From tubulated endo-lysosomes reformed LYSs emerge (Step 3). ‘Secretory’ LEs translocate to the cell periphery (Step 4) and release their sterol content into the extracellular medium via transient fusion with the PM as exosomes (light-blue rings; Step 5). Sterol release could also take place by non-vesicular sterol transfer from peripheral LE/LYSs to the PM, possibly via ER-PM contact zones (Step 6). The latter process could be followed by extracellular shedding of micro-vesicles to relieve the sterol load in the PM. The ‘secretory’ LEs are characterized by their fusion competence to receive cargo from early endosomes (Step 7). See text for further discussion.

In summary, our multi-modal optical imaging study allows us to suggest a model for sterol export from LE/LYSs by NPC2 (Fig. 9). We propose that NPC2 selectively enhances non-vesicular sterol export from LE/LYSs by supplying sterol from ILVs to the limiting membrane of LE/LYSs for further export via NPC1 or other endo-lysosomal membrane proteins ^30^. Clearance of autophagosomes, active centripetal transport of LE/LYSs and restoration of their fusion competence are suggested to be consequences of this NPC2 mediated selective sterol export. Shedding of ectosomes from the PM as well as stimulated release of ILVs from multi-vesicular endosomes are likely mechanisms for the final steps in sterol efflux. Further studies are warranted to determine the molecular players, which orchestrate sterol flux through endo-lysosomes together with NPC2 in living cells.

## Materials and Methods

### Reagents

Fetal bovine serum and DMEM were from GIBCO BRL (Life Technologies, Paisley, Scotland). Other chemicals including human lipoprotein depleted serum (LPDS) and DHE were from SIGMA Chemical (St. Louis, MO). Rhodamine-labeled dextran (Rh-dextran; 70kD), succinimidyl esters of Alexa488, Alexa546 and Alexa647 as well as C6-nitrobenzoxadiazole (NBD)-Ceramide (C6-NBD-Cer) and Alexa488-tagged bovine serum albumin (Alexa488-BSA) were purchased from Invitrogen/Molecular Probes (Inc. USA). Buffer medium contained 150 mM NaCl, 5 mM KCl, 1 mM CaCl_2_, 1 mM MgCl_2_, 5 mM glucose and 20 mM HEPES (pH 7.4) as described ^73^. Transferrin (Tf) was iron loaded as previously described ^74^. Succinimidyl ester of Alexa647 dye (emission in infrared) was then conjugated to the iron-loaded Tf following the manufacturer’s instructions. NPC2 was purified from bovine milk and conjugated with succimidyl esters of Alexa488 (emission in green) or of Alexa546 dye (emission in red) as described previously ^21^.

#### Cell culture

NPC2 deficient human skin fibroblasts were from Coriell Institute #GM18455 (a male patient affected by two point mutations at the NPC2 locus resulting in a nonsense mutation at codon 20 in allele 1 and in a missense mutation at codon 47 of allele 2, respectively), while human skin fibroblasts from control subjects (Coriell Institute #GM08680) were from a male healthy donor ^30^. Both cell types were grown at 37°C in an atmosphere of 5% CO_2_ until 90% confluence in complete DMEM culture medium supplemented with 1% glutamine, 1% penicillin and 20% FBS for diseased cells or 10% FBS for control cells. The cells were placed on microscopy dishes and allowed to settle for another 48 hours before the experiments.

#### Sterol efflux experiments

##### Sterol efflux under continuous presence of fluorescent NPC2

Cells were incubated with the DHE/BSA solution in medium containing LPDS containing for 48 h, washed and incubated with 100 nM Alexa488-NPC2 for 8-96h in medium with LPDS (Protocol A, Fig. S1). During the last 4h, 0.5 mg/ml Rh-dextran and during the last 30 min of incubation additionally 5 μg/ml Alexa647-Tf were added to the medium. Cells were subsequently washed and imaged as described below. In parallel experiments, cells were after labeling with DHE from a DHE/BSA complex incubated in culture medium with LPDS but no NPC2 (Protocol C, Fig. S1). Cells were washed and imaged in buffer medium, as described below.

##### Sterol efflux after removal of fluorescent NPC2 from the medium

Cells were incubated with the DHE/BSA solution in medium containing LPDS containing for 24 h, washed and incubated with 200 nM Alexa488-NPC2 for 24 h in medium with LPDS, washed and incubated for up to 72h in medium with LPDS (Protocol B, Fig. S1).

### Fluorescence microscopy

Wide field epifluorescence microscopy was carried out on a Leica DMIRBE microscope with a 63 x 1.4 NA oil immersion objective (Leica Lasertechnik GmbH) controlled by a Lambda SC smart shutter (Sutter Instrument Company). Images were acquired with an Andor Ixon^EM^ blue EMCCD camera driven by the Solis software supplied with the camera. DHE was imaged in the UV using a specially designed filter cube obtained from Chroma Technology Corp. with 335-nm (20-nm bandpass) excitation filter, 365-nm dichromatic mirror and 405-nm (40-nm bandpass) emission filter. Rh-dextran and Alexa546-NPC2 were imaged using a standard rhodamine filter set [535-nm, (50-nm bandpass) excitation filter, 565-nm longpass dichromatic filter and 610-nm (75-nm bandpass) emission filter], while Alexa488-NPC2 was imaged using a standard fluorescein filter set [470-nm, (20-nm bandpass) excitation filter, 510-nm longpass dichromatic filter and 537-nm (23-nm bandpass) emission filter]. Alexa647-Tf was detected with an infrared filter cube [620-nm, (20-nm bandpass) excitation filter, 660-nm dichromatic mirror, and 700 nm (75-nm bandpass) emission filter]. For detecting co-localization of DHE with organelle markers, a correction for chromatic aberration was performed as described ^75^.

#### Fluorescence recovery after photobleaching (FRAP) experiments

Cells were labeled with DHE/BSA for 24 h as described above, washed and incubated with 100 nM Alexa546-NPC2 for another 24h, washed and placed on the stage of the wide field microscope. Images were taken in the UV channel (the DHE prebleach image) and in the red channel (for Alexa546-NPC2) by taking chromatic aberration into account, as described ^75, 76^. The bleached region was determined by closing the field aperture in the microscope optical train thereby narrowing the field of view to a circular region of radius 15 μm. Another reference image was taken in the red channel to determine position and size of the region of interest (ROI) for the bleach before the same ROI was bleached in the DHE channel for five sec by continuous illumination from the mercury arc lamp ^76^.

Subsequently the field aperture was opened, a 50% neutral density filter was inserted and images were acquired in the DHE channel every 20 sec. A short acquisition time (typically 5-10 msec) was chosen to minimize photobleaching during recording of fluorescence recovery sequences. Based on the red channel image of Rh-dextran, the focus was set to the LE/LYSs while taking the chromatic shift between the UV-and red channel into account. For FRAP in the PM, the ROI was moved to the cell periphery to avoid bleaching underlying organelles.

### Image analysis of fluorescence microscopy data

#### Quantification of sterol efflux and sorting from internalized fluorescent NPC2s

For quantification of cellular DHE or Alexa546-NPC2 by digital image analysis, images were always first background-corrected as described using either MatLab (The MathWorks) or the open-source image analysis software ImageJ (developed at the U.S. National Institutes of Health and available on the Internet at http://rsb.info.nih.gov/ij) ^77-79^. Before quantification, the digital (i.e. pixel) resolution of the images was enhanced using an image interpolation routine in MatLab. More precisely, images features such as edges were enhanced with a sharpening filter. Then the image pixel size was increased by a cubic interpolation.. This made it easier and more reliable to detect and segment particle-like structures in the images, especially in the case of DHE (see Supplementary Information, section S2). Image segmentation for intensity quantification per vesicle was performed by particle detection followed by seed-based watershed segmentation using self-written or pre-assembled routines in MatLab. Distance-to-membrane calculations, co-localization and RDF analyses were performed using SpatTrack ^21^.

#### Quantification of fluorescence recovery of DHE

A macro was written within ImageJ, which automatically performs a background subtraction, measures the integrated intensity in the bleached area and in the whole cell, as outlined manually, and calculates the ratio of DHE fluorescence in the bleached region divided by the total cellular DHE intensity, as described previously ^30^. This procedure efficiently accounted for any photobleaching during image acquisition, as we showed in experiments ^76^ and simulations ^80^. Subsequently, the resulting fractional recovery was normalized to the initial postbleach intensity (i.e., the first data point in the recovery curve) and fitted to a mono-exponential recovery function for each vesicle separately in ImageJ. For estimation of the intensity profile across the ROI during recovery, the Radial profile plot plugin developed by Paul Baggethun was used in ImageJ (https://imagej.nih.gov/ij/plugins/radial-profile.html). For visualizing the recovery, as shown in Fig. S11 in the Supplementary Information section, our previously developed ImageJ Macro was used which measures the pixel-wise intensity and normalizes each image to the average intensity in the pre-bleach image ^30^. This macro generates a new stack in which the recovering DHE fluorescence in the bleached ROI was not obscured by photobleaching during image acquisition and is used for visualization purposes but not for quantification.

#### Object-based co-localization and spatial pattern analysis of LE/LYS

Co-localization between DHE and Alexa546-NPC2 was calculated in an object-based manner using the PDBCA subroutine in SpatTrack, as recently described ^21^. The radial distribution function (RDF) was calculated for DHE-containing LE/LYSs in direct comparison to a random particle distribution, exactly as recently described for LE/LYSs containing Alexa546-NPC2 ^21^. In addition, we calculated the ratio of co-localized vesicles to all vesicles of that type (i.e., DHE or Alexa546-NPC2) as a function of distance to the membrane. That is, for each distance increment the number of e.g. DHE containing vesicles being co-localized with NPC2 containing vesicles is normalized with respect to the total number of DHE containing vesicles at that distance. Since there are different numbers of DHE and NPC2 containing vesicles as a function of distance to the PM, this analysis was performed for both types of vesicles.

##### Cell preparation for soft X-ray tomography

Healthy and NPC2 deficient human fibroblasts were grown to a confluency of 90%, trypsinized and split onto Poly-D-Lysine coated R 2/2 grids (QUANTIFOIL, 100 Holey Carbon Films, Grids: HZB-2 Au) fixed to the bottom of 12 well plates. Cells were allowed to settle for another 48-72 h before further treatment and fixed with 4% PFA. Prior to fixation NPC2 lacking fibroblasts were incubated with 100 nM or 200 nM NPC2-Alexa546 protein in media containing LPDS for 72 h and healthy fibroblasts were incubated with 0.5 mg/mL Rh-dextran in complete growth medium overnight, at 37°C in an atmosphere of 5% CO2.

The cells were kept in PBS until cryo-plunge freezing with liquid ethane and subsequently storage in liquid nitrogen. The imaging was performed at liquid nitrogen temperature to prevent recrystallization of the frozen liquid. Prior to the plunge freezing a small volume of ~270 nm gold shelf beads were added to the samples, to serve as fiducial markers for tomographic reconstruction.

##### Soft X-ray tomography

SXT was performed at the electron storage ring BESSY II operated by the Helmholtz-Zentrum Berlin. The SXT data was collected at the full-field transmission X-ray microscope (TXM), with an x-ray energy of 510 eV and a 25 nm zone plate. Cells were kept at liquid nitrogen temperature during imaging, and imaged over a tilt range of 120-125° with 1° step size. The image pixel size was 9.8 nm. For collecting the corresponding fluorescence signal, a widefield microscope connected to the X-ray microscope, with an Zeiss LD EC Epiplan Neofluar 100/0.75 DIC was used ^81^.

#### SXT image processing and analysis

The B-soft^82^ and Tomo3D^83^ software were used to align SXT projections and to reconstruct the three-dimensional SXT data, respectively. The segmentation and subsequent rendering were performed in the free available software Ilastik (http://ilastik.org/index.html) ^84^. Before segmentation, the tomography images were pre-processed in ImageJ with a median filter. Montages, sum z-projections and measurements of vesicle sizes were all performed with the use of ImageJ. The statistics software package R (http://www.r-project.org) was used for further analysis of vesicle sizes. The datasets of the endosome sizes in NPC2-/- cells incubated without NPC2 (n=134 vesicles, N=3 cells), in NPC2-/- cells incubated for 72 h with NPC2 (n=120, N=3) and in control fibroblasts (n=86, N=1) were transformed to log normal distributions for which ANOVA and Tukey HSD tests were performed. The artistic reconstruction of observed features was created in Microsoft Paint 3D.

## Acknowledgements

Financial support from the Novo Nordisk and Villum foundations to DW, Denmark is gratefully acknowledged. This work was also supported by the VILLUM Center for Bioanalytical Sciences at the University of Southern Denmark. We thank HZB for the allocation of synchrotron radiation beamtime at the undulator beamline U41-TXM at the BESSY II electron storage ring, Berlin.

## Author contributions

AD, MLVJ and MS carried out fluorescence experiments, AD, MS, PG, SW, JM, DW and GS carried out SXT experiments and provided SXT instrumentation, respectively. GKN and CWH purified and provided NPC2 protein. AD, FWL and DW analyzed fluorescence data, SK, AD and DW analyzed SXT data, DW wrote the manuscript with help and feedback from the other authors.

## Data availability statement

Experimental microscopy data and program code for image analysis routines will be made available by the authors upon request.

## Appendix Kinetic analysis of sterol efflux from LE/LYSs using a stochastic compartment model

Sterol efflux is modelled by a stochastic irreversible continuous-time one-compartment model with time-dependent hazard rate. The survival probability is a measure for the likelihood of sterol molecules to ‘survive’ in the LE/LYSs until time, *t*. Mathematically, it reads for the Weibull distribution ^33^:

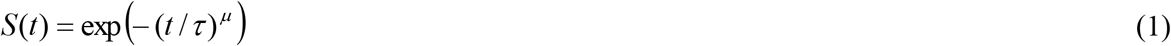

Here, τ is the time constant of the decay and μ is the stretching exponent describing the heterogeneity of the observed process. To fit this model to our data, we multiply with the initial sterol amount in internal compartments (i.e., mostly LE/LYSs and some early and recycling endosomes), called *S*_0_, and add the sterol fraction in LE/LYSs of healthy fibroblasts as background term, *B*, in the fitting procedure. Thus, we fit the experimental data to:

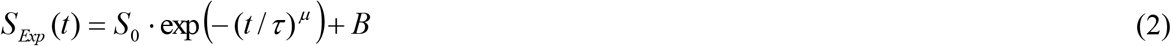

The *pdf* of this Weibull model is a measure for the likelihood of sterol to leave the cells and reads:

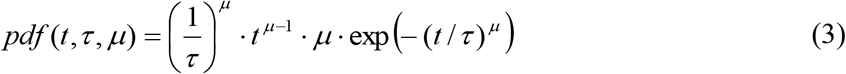

The rate coefficient also called a hazard rate is the negative of the time-derivative of the logarithm of the survival probability ^33^. It is given for the Weibull distribution by:

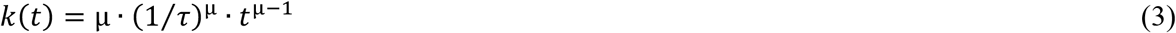

It is plotted as function of time for the estimated parameter values in Fig. S2.

